# Environmental determinants of pyoverdine production, exploitation and competition in natural *Pseudomonas* communities

**DOI:** 10.1101/263004

**Authors:** Elena Butaitė, Jos Kramer, Stefan Wyder, Rolf Kümmerli

## Abstract

Many bacteria rely on the secretion of siderophores to scavenge iron from the environment. Laboratory studies revealed that abiotic and biotic factors together determine how much siderophores bacteria make, and whether siderophores can be exploited by non-producing cheaters or be deployed by producers to inhibit competitors. Here, we explore whether these insights apply to natural communities, by comparing the production of the siderophore pyoverdine among 930 *Pseudomonas* strains from 48 soil and pond communities. We found that pH, iron content, carbon concentration, and community diversity determine pyoverdine production levels, and the extent to which strains are either stimulated or inhibited by heterologous (non-self) pyoverdines. While pyoverdine non-producers occurred in both habitats, their prevalence was higher in soils. Environmental and genetic analysis suggest that non-producers can evolve as cheaters, exploiting heterologous pyoverdine, but also due to pyoverdine disuse in environments with increased iron availability. Overall, we found that environmental factors explained between-strain variation in pyoverdine production much better in soils than in ponds, presumably because high strain mixing in ponds prevents local adaption. Our study sheds light on the complexity of natural bacterial communities, and provides first insights into the multivariate nature of siderophore-based iron acquisition and competition among environmental pseudomonads.

## Introduction

Iron is a key growth-limiting factor for most bacteria. It is required for various fundamental cellular processes, such as DNA biosynthesis and respiration (Andrews *et al*., 2003). However, iron is often limited in nature because it is either insoluble in its ferric (Fe^+3^) form at circumneutral pH and aerobic conditions, host-bound in the context of infections, or occurs only at very low concentrations in habitats like the ocean (Andrews *et al*., 2003; Miethke and Marahiel, 2007; Boyd and Ellwood, 2010). To overcome iron limitation, many bacteria secrete siderophores, iron-scavenging molecules that have a high affinity for ferric iron (Neilands, 1981; Hider and Kong, 2010). These molecules bind iron from natural sources and can then be taken up by cells via specific receptors. Given the importance of iron, it comes as no surprise that bacteria have evolved sophisticated mechanisms to accurately adjust the level of siderophore production to match prevailing levels of iron limitation (Andrews *et al*., 2003; Parrow *et al*., 2013; Frawley and Fang, 2014).

In addition to their role in provisioning producers with iron, siderophores can also have fitness consequences for other community members, including non-producers and individuals with different siderophore systems (Loper and Henkels, 1999; D’Onofrio *et al*., 2010; Cordero *et al*., 2012; Traxler *et al*., 2012, Galet *et al*., 2015; Bruce *et al*., 2017; Butaitė *et al*., 2017). This is because siderophores can be shared between individuals with compatible receptors, which can select for exploitation by cheating strains that do not produce or produce less of the specific siderophore, yet capitalize on the siderophores produced by others (West *et al*., 2006; Jiricny *et al*., 2010). Moreover, siderophores can lock iron away from competitors with incompatible receptors, and can therefore be involved in inter-strain competition (Buyer and Leong, 1986; Joshi *et al*., 2006; Butaitė *et al*., 2017; Niehus *et al*., 2017; Schiessl *et al*., 2017; Sexton *et al*., 2017; Leinweber *et al*., 2018). Although laboratory studies have uncovered many of the molecular, ecological and evolutionary aspects of fine-tuned siderophore regulation and siderophore-mediated social interactions, we know surprisingly little about the determinants of siderophore production under natural conditions in complex multi-species communities.

Here, we address this issue by examining how environmental factors of natural soil and pond habitats relate to the amount of siderophores produced by their inhabitants, and the ability of natural isolates to affect other members of their community via their siderophores. In our study, we focus on the phylogenetically diverse group of fluorescent pseudomonads, whose members are known to produce the green-fluorescent pyoverdine as their primary siderophore (Meyer *et al*., 2008; Cornelis, 2010; Cézard *et al*., 2015). From laboratory studies, predominantly carried out with *Pseudomonas aeruginosa*, we know that iron concentration, organic carbon composition, pH, and community diversity all influence the level of pyoverdine production. Specifically, bacteria gradually downscale pyoverdine production in response to increased iron concentration (Tiburzi *et al*., 2008; Kümmerli *et al*., 2009), when iron is bound to relatively weak organic chelators (Dumas *et al*., 2013), and at low pH where solubility of iron is increased (Loper and Henkels, 1997). In addition, *P. aeruginosa* upregulates pyoverdine production in the presence of competing strains or species (Harrison *et al*., 2008; Kümmerli *et al*., 2009). Interestingly, pyoverdine-mediated social interactions were found to be influenced by similar environmental factors. For example, increased iron availability allows producers to down-scale pyoverdine production, which lowers their susceptibility to exploitation by non-producers (Kümmerli *et al*., 2009). Similarly, increased carbon availability reduces the relative metabolic costs of pyoverdine production, and thus decreases the advantage of cheaters (Brockhurst *et al*., 2008; Sexton and Schuster, 2017). Finally, community composition was found to be a key factor determining who interacts with whom, thereby influencing the relative importance of competition versus cooperation (Griffin *et al*., 2004; Inglis *et al*., 2016; Leinweber *et al*., 2017). To examine whether these factors influence pyoverdine production and pyoverdine-mediated social interactions in natural communities, we isolated a total of 930 pseudomonads from 24 soil cores and 24 pond water samples. We measured the pyoverdine production of all isolates in standard iron-limited medium, and related this measure to the total iron and carbon concentrations as well as the pH of their environment, and to the phylogenetic diversity of their community. For a subset of strains, we additionally quantified the effect of pyoverdine-containing supernatant on the fitness of co-occurring strains. For these experiments, we focused on pyoverdine non-producers as supernatant-recipients because the stimulation or inhibition of their growth under iron-limited conditions provides direct evidence for whether or not they possess matching receptors for the uptake of heterologous (non-self-produced) siderophores (Butaitė *et al*., 2017).

## Results

### Pyoverdine production profiles differ between soil and pond pseudomonads

Prior to phenotypic screening, we sequenced the housekeeping *rpoD* gene, commonly used for phylogenetic affiliation of pseudomonads, for all 930 isolates to confirm that they indeed belong to this taxonomic group (Fig. S1). Once this was confirmed, we measured the pyoverdine production of all isolates (using the natural fluorescence of this molecule) in iron-limited casamino acids (CAA) medium supplemented with transferrin as an iron chelator. We found that isolates greatly varied in the amount of pyoverdine they make (Fig. 1A). While some strains produced no measurable amount of pyoverdine, others produced much more than our laboratory reference strains (pyoverdine values are scaled relative to the average production level of the laboratory reference strains listed in Table S1). Across all isolates, there was a positive correlation between overall pyoverdine levels and their growth in iron-limited CAA, suggesting that pyoverdine is important to overcome iron limitation (linear mixed model, LMM: *t*_917.7_ = 18.62, *p* < 0.001; Fig. 1B).

**Figure 1.**
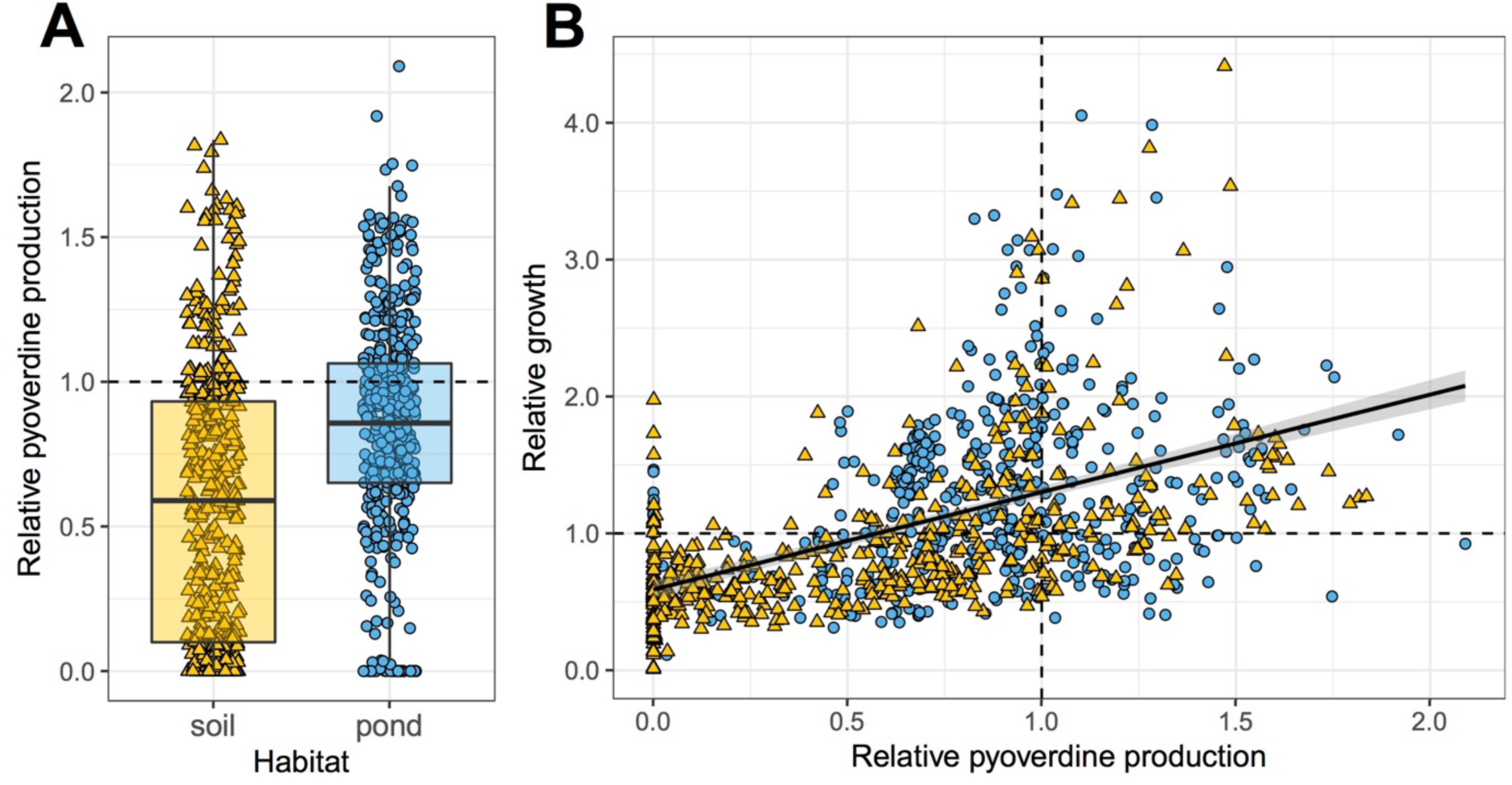
Environmental pseudomonads vary greatly in their level of pyoverdine production and ability to grow under iron-limited conditions. **(A)** Our phenotypic screen involving 930 natural pseudomonads from 24 soil and 24 pond communities (*n* = 462 soil and *n* = 468 pond) revealed that soil isolates produce significantly less pyoverdine than pond isolates, and that there are significantly more non-producers in soil than in pond communities. **(B)** There is a positive correlation between the growth (optical density measured at 600 nm) of isolates and their pyoverdine production level, indicating that pyoverdine is important to cope with iron limitation. Pyoverdine production and growth were measured in iron-limited CAA medium, and scaled relative to the values of eight laboratory reference strains (Table S1). Values represent means across three replicates for soil (yellow triangles) and pond (blue circles) isolates. Dashed lines indicate pyoverdine production (horizontal line in A, vertical line in B) and growth (horizontal line in B) of the reference strains. The box plot shows the median (bold line), the 1^st^ and 3^rd^ quartile (box), and the 5^th^ and 95^th^ percentile (whiskers).

There were several significant differences between soil and pond isolates. First, the relative pyoverdine production was significantly lower for soil than for pond isolates (mean ± SE for soil isolates: 0.571 ± 0.022; for pond isolates: 0.832 ± 0.018; LMM: *t*_21.5_ = −4.36, p < 0.001; Fig. 1A). Second, there were significantly more pyoverdine non-producers (defined as producing less than 5% of laboratory reference strains) in soil than in pond communities (19.7% vs. 8.3%; Fisher’s exact test: *p* < 0.0001; maximal non-producer frequency per community was 75% and 20% in soils and ponds, respectively). Finally, the coefficient of variation (CV) in relative pyoverdine production among isolates from the same community was significantly higher in soil than in pond communities (mean CV ± SE for soil communities: 85.2% ± 8.3%; for pond communities: 46.2% ± 2.4%; LMM: *t*_22.0_ = 4.39, *p* < 0.001; Fig. S2).

To explore the relationship between phylogenetic diversity and pyoverdine production, we constructed maximum likelihood phylogenetic trees based on partial *rpoD* gene sequences for soil and pond isolates (Fig. S1). We found that phylogenetic diversity (normalized by the number of isolates per community) was significantly lower in soil than pond communities (Faith’s phylogenetic diversity, median ± [1st quartile | 3rd quartile], for soil communities: 0.09 ± [0.07 | 0.11]; for pond communities: 0.11 ± [0.09 | 0.15]; Mann-Whitney U-test: *W* = 178, *p* = 0.023).

We further examined whether there is a phylogenetic signal for pyoverdine production (i.e. whether closely related isolates show similar pyoverdine production) (Fig. S1). In line with our previous findings, based on a smaller dataset (Butaitė *et al*., 2017), we found a very weak phylogenetic signal for soil communities (Blomberg’s *K* = 0.02 ± [0.01 | 0.12]), and a low but slightly increased phylogenetic signal in pond communities (*K* = 0.12 ± [0.03 | 0.25]; test for differences between soil and pond communities, Wilcoxon rank sum test: *W* = 186, *p* = 0.036). These analyses confirmed that pyoverdine production levels can vary substantially even among closely related strains, and thus suggest that pyoverdine non-producers evolved independently on multiple occasions.

### Environmental determinants of pyoverdine production

Next, we tested for a relationship between the pyoverdine production profiles described above and different abiotic (pH, total iron and carbon concentrations) and biotic (phylogenetic community diversity) variables of the environments the isolates originated from. These environmental variables varied considerably between communities (Fig. S3, Table S2), and were often correlated across communities (Table S3). To account for the resulting collinearities, we first carried out separate principal component analyses (PCAs) for soil and pond that each incorporated the four environmental variables. PCAs break collinearities by converting the values of potentially correlated variables into a set of uncorrelated variables called principal components (PCs). Here, we focused on the first two PCs (henceforth called SPC1 and SPC2 for soil, and PPC1 and PPC2 for pond), which explained 88.0% (soil) and 88.3% (pond) of the total variance observed (Table 1). Each PC is characterized by a set of positive and/or negative loadings, which describe how much it is influenced by each of the environmental variables fed into the PCA. These PCs were then used in standard linear models to test whether they correlate with the pyoverdine production levels of our isolates.

**Table 1.** Loadings of the abiotic and biotic environment-defining variables onto the principal components (PCs) for **(A)** soil and **(B)** pond communities.

| **(A) Soil** | **SPC1** | **SPC2** | SPC3 | SPC4 |
| --- | --- | --- | --- | --- |
| community diversity | -0.1729 | **0.9358** | -0.2445 | -0.1862 |
| pH | **-0.5396** | 0.1584 | **0.7812** | 0.2711 |
| iron | **-0.5751** | -0.1406 | -0.5674 | **0.5723** |
| carbon | **0.5901** | 0.2819 | 0.090 | **0.7512** |
| explained variance [%] | 62.3 | 25.7 | 9.9 | 2.1 |

| **(B) Pond** | **PPC1** | **PPC2** | PPC3 | PPC4 |
| --- | --- | --- | --- | --- |
| community diversity | **-0.4892** | -0.3813 | **-0.7797** | 0.0857 |
| pH | **-0.5161** | **0.5353** | -0.0114 | **-0.6686** |
| iron | **0.5449** | -0.3705 | -0.2391 | **-0.7132** |
| carbon | **0.4443** | **0.6564** | **-0.5786** | 0.1924 |
| explained variance [%] | 69.5 | 18.8 | 9.2 | 2.5 |

SPC = Soil Principal Components
PPC = Pond Principal Components
Following Jolliffe (1972), we retained those PCs (in bold) with an eigenvalue greater than 0.7 times the average eigenvalue. Similarly, we considered the loading of a variable onto a PC to be significant (in bold) if the loading was greater than 0.7 times the highest loading of the PC.

For soil communities, SPC1 was positively loaded by carbon concentration, and negatively by pH and iron concentration, indicating that SPC1 reflects a trade-off (i.e., a negative correlation) between carbon concentration on the one hand, and pH and iron concentration on the other hand. By contrast, SPC2 was solely and positively loaded by community diversity (Table 1A). When relating these PCs to the pyoverdine production profiles of the soil isolates, we found a positive correlation between relative pyoverdine production and community diversity (SPC2; *t*_18.3_ = 3.36, *p* = 0.0034; Fig. 2A). In contrast, there was no association between the relative pyoverdine production and the trade-off between carbon versus pH and iron captured by SPC1 (*t*_10.6_ = −1.93, *p* = 0.0809). Because there were many pyoverdine non-producers, present in 22 out of 24 soil communities, we further examined whether the likelihood of being a pyoverdine non-producer correlated with the SPCs. We found that the likelihood of being a non-producer was highest in communities characterized by relatively high levels of carbon combined with relatively low levels of pH and iron (SPC1; *z* = 2.85, *p* = 0.0044; Fig. 2B). Conversely, the likelihood of being a non-producer did not correlate with community diversity (SPC2; *z* = −0.72, *p* = 0.4745).

**Figure 2.**
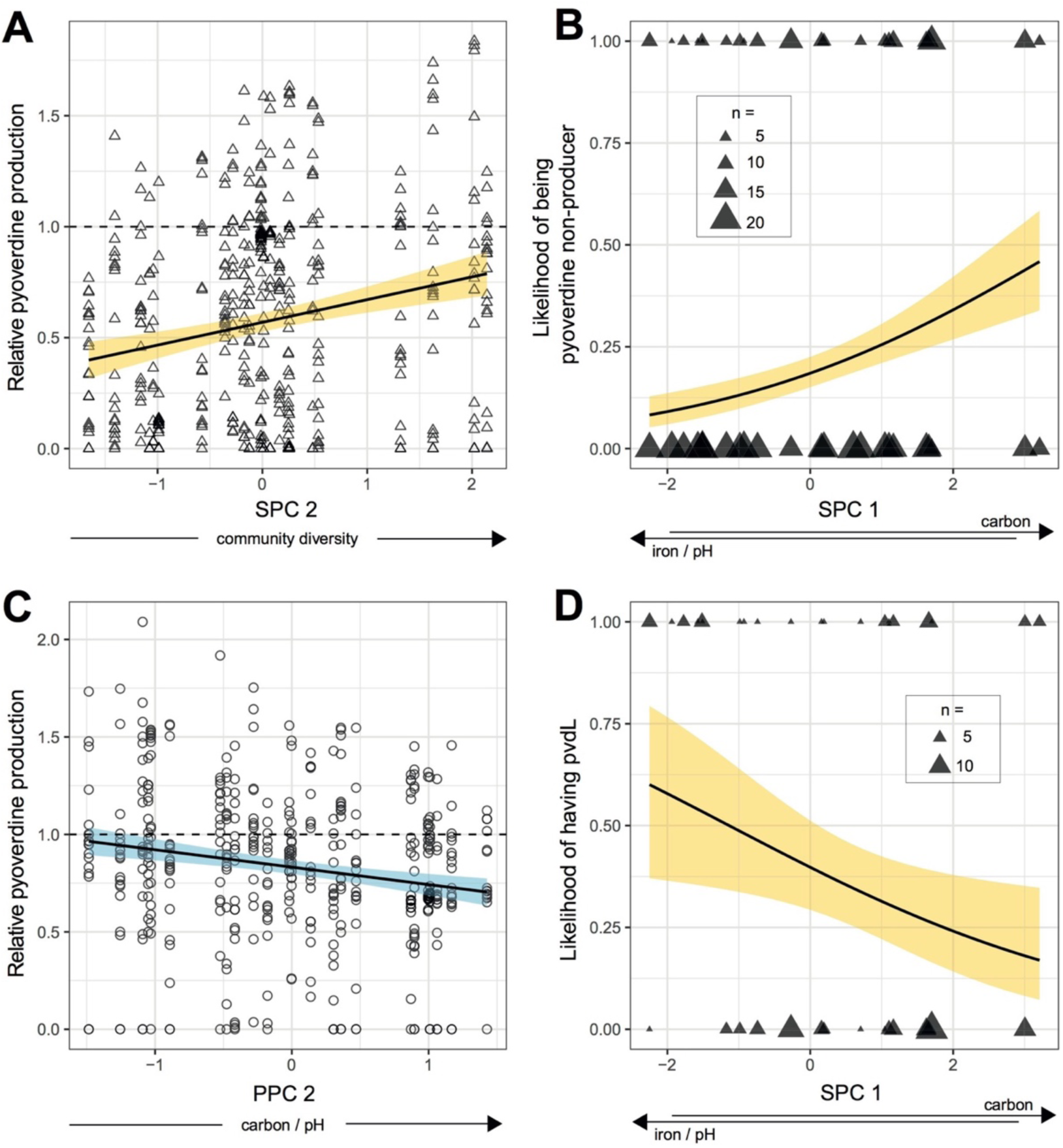
Correlations between parameters of pyoverdine production and environmental variables. In soil communities (*n* = 462 isolates), relative pyoverdine production in iron-limited medium increased with community diversity (SPC2) **(A)**, whereas the likelihood of being a pyoverdine non-producer correlated with SPC1, i.e. increased with higher concentration of carbon in combination with lower pH and reduced iron concentration **(B)**. For pond communities (*n* = 468 isolates), relative pyoverdine production correlated negatively with PPC2, i.e. decreased with higher concentrations of of carbon in combination with increased pH **(C)**. A PCR amplification of the gene *pvdL* in the 91 non-producers from soil revealed a negative correlation between the likelihood of having *pvdL* and SPC1 **(D)**. Since *pvdL* serves as an indicator for the presence of the pyoverdine biosynthesis locus, this analysis indicates that the likelihood of still having *pvdL*, although being a non-producer, increases in habitats characterized by high pH and iron in combination with low carbon. Solid lines with shaded areas show significant correlations together with the 95% confidence band for soil (yellow) and pond (blue) communities. Dashed lines depict scaled pyoverdine production of reference strains (Table S1).

For pond communities, PPC1 was positively loaded by iron and carbon, and negatively by pH and community diversity, indicating that PPC1 reflects a trade-off between these groups of environmental variables (Table 1B). Conversely, PPC2 was positively loaded by pH and carbon. Note that this positive association is independent of and occurs simultaneously with the trade-off between these variables in PPC1 (Table 1B). When feeding these PCs into a linear mixed model we found a negative association between relative pyoverdine production and PPC2 (*t*_10.9_ = −2.92, *p* = 0.0140), indicating that the relative pyoverdine production was lower among pond isolates from communities characterized by higher levels of carbon and pH (Fig. 2C). In contrast, PPC1 was not linked to the pyoverdine production of pond isolates (*t*_10.0_ = 0.68, *p* = 0.5120). We further examined whether the likelihood of being a pyoverdine non-producer correlated with one of the PPCs, but this was neither the case for PPC1 (*z* = −1.16, *p* = 0.2450) nor for PPC2 (*z* = 0.79, *p* = 0.4320).

### The genetic basis of pyoverdine non-production

The occurrence of many pyoverdine non-producers prompted us to explore their genetic makeup in more detail. Our previous study (Butaitė *et al*., 2017), entailing whole-genome sequencing of environmental isolates, revealed two different types of non-producers: those with a highly truncated pyoverdine locus, and those with a seemingly intact, yet silent locus. To find out whether these types also occur here, and whether the frequency of the two types correlates with environmental variables, we screened all 130 non-producers for the presence of the *pvdL* gene. This gene encodes an essential and conserved non-ribosomal peptide synthetase involved in pyoverdine synthesis (Ravel and Cornelis, 2003; Smith *et al*., 2005). The presence or absence of the *pvdL* gene hence indicates whether non-producers possess a silent yet intact locus or a truncated dysfunctional locus, respectively.

We found that 32 out of 91 (35.2%) soil non-producers and 15 out of 39 (38.5%) pond non-producers were positive for the presence of *pvdL* (no significant difference between soil and pond: *χ*^2^ = 2.48, *p* = 0.1151). For soil communities, the likelihood of being *pvdL*-positive was lowest in communities characterized by a combination of relatively high carbon concentration, and a low pH and iron concentration (SPC1: *z* = −2.07, *p* = 0.0386; Fig. 2D). For pond communities, meanwhile, none of the two PPCs were associated with the likelihood of being *pvdL*-positive (PPC1: *z* = −1.54, *p* = 0.1230; PPC2: *z* = −0.83, *p* = 0.4090).

### Supernatants from pyoverdine producers affect the growth of non-producers

To estimate the extent to which pyoverdine could be involved in social interactions between strains, we compared the growth of non-producers in iron-limited medium alone versus their growth in medium supplemented with pyoverdine-containing supernatants of producers from the same community. Overall, we fed 53 non-producers (from 12 soil and pond communities each) with pyoverdine-containing supernatants from four to six different producers from the same community. This resulted in a total of 152 soil and 151 pond non-producer-supernatant combinations.

We found that the fitness effects of pyoverdine-containing supernatants on non-producers covered the entire range from almost complete growth inhibition to high stimulation (Fig. S4). While the likelihood of stimulation did not significantly differ between the two habitats (soil vs. pond: *z* = 1.80, *p* = 0.072; Fig. 3A), the likelihood of inhibition was significantly higher in pond than in soil communities (*z* = 2.86, *p* = 0.004; Fig. 3B). Note that ‘stimulation’ or ‘inhibition’ describe the respective cases where non-producers grew significantly better or worse, with pyoverdine-containing supernatants than without. Although supernatants contain other growth-modulating components in addition to pyoverdine, we have previously shown that the growth effects of this supernatant assay on non-producers are mainly caused by pyoverdine, and are likely driven by the presence (stimulation) and absence (inhibition) of a matching receptor for pyoverdine uptake (Butaitė *et al*., 2017).

**Figure 3.**
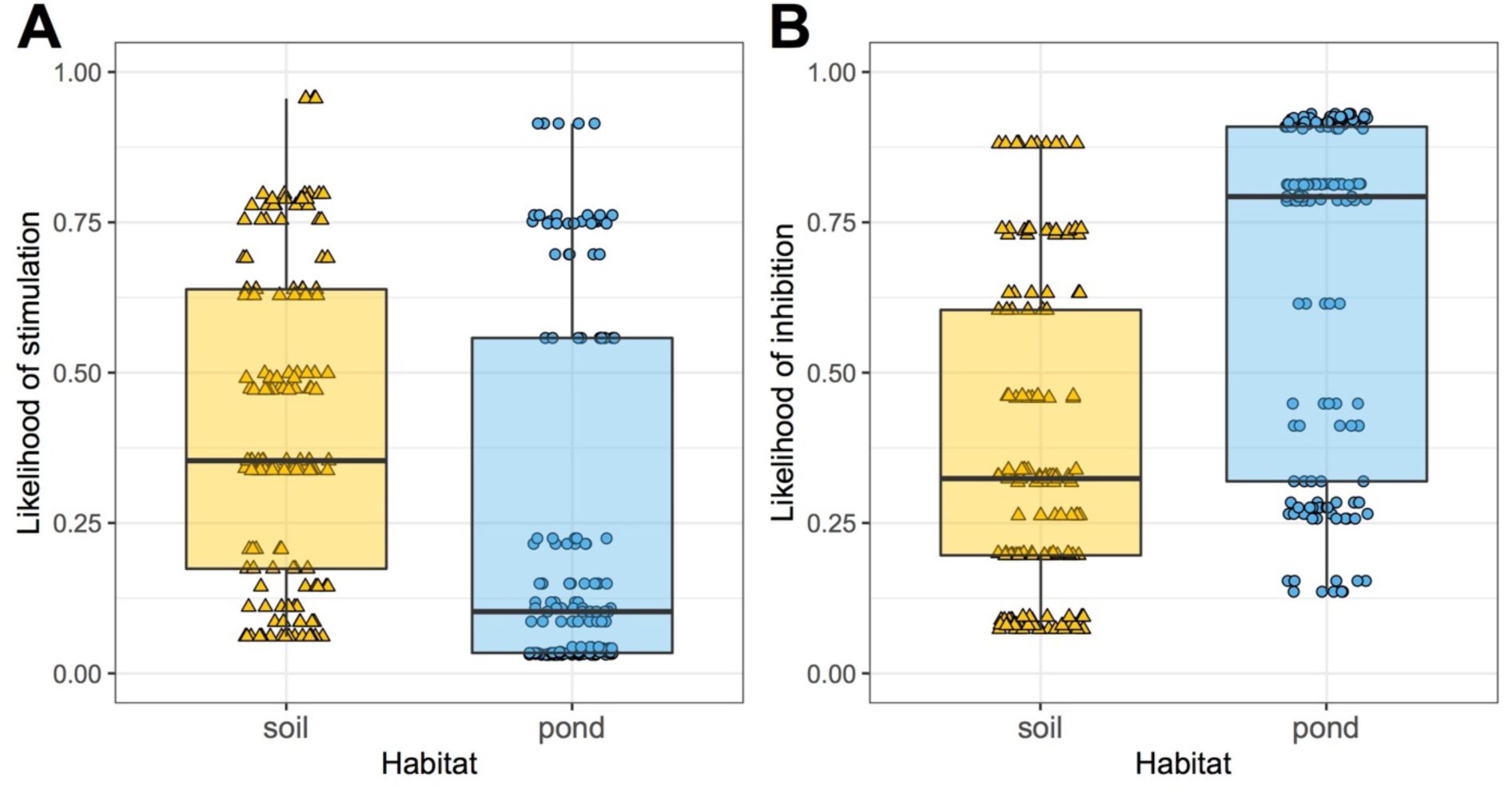
Pyoverdine-containing supernatants from producers can both stimulate and inhibit the growth of non-producers. **(A)** The likelihood for non-producers to be stimulated by pyoverdine-containing supernatants from a producer did not significantly differ between pond and soil isolates. **(B)** In contrast, the likelihood for non-producers to be inhibited by producer supernatants was significantly higher for pond than for soil isolates. The supernatant assay involved 53 (soil 26 / pond 27) non-producers, and 142 (70/72) producer supernatants from a subset of communities (12/12), resulting in a total of 303 (152/151) non-producer-supernatant combinations, each replicated three to four times. ‘Stimulation’ or ‘inhibition’ corresponded to cases where non-producers grew significantly better or worse with supernatant than without, respectively. Box plots show the median (bold line), the 1^st^ and 3^rd^ quartile (box), and the 5^th^ and 95^th^ percentile (whiskers).

### Environmental determinants of pyoverdine-mediated social interactions

Next, we tested whether the likelihood of stimulation and inhibition correlate with environmental factors. To this end, we reused the first two PCs describing the relationships between our four environmental variables of interest (pH, iron and carbon concentrations, community diversity) as explanatory variables. Because the supernatant assay involved pairs of strains (i.e. the supernatant donor and the recipient), we further included the phylogenetic relatedness between isolates, based on *rpoD* sequence similarity, into our statistical models. For soil communities, we found that the likelihood of stimulation correlated with SPC1, suggesting that non-producers were most likely to be stimulated by heterologous pyoverdine when originating from environments featuring a combination of a high carbon concentration on the one hand, and a low pH and iron concentration on the other hand (*z* = 2.21, *p* = 0.0269; Fig. 4A). Furthermore, the likelihood of stimulation tended to increase with community diversity (SPC2; *z* = 1.96, *p* = 0.0502), and increased with the relatedness between strains (*rpoD* identity; *z* = 3.44, *p* = 0.0006; Fig. 4B). The opposite trends were generally observed for the likelihood of inhibition: it correlated negatively with SPC1 (*z* = −1.94, *p* = 0.0525), and relatedness (*rpoD* identity; *z* = −2.80, *p* = 0.0052), but not with community diversity (SPC2; *z* = −1.01, *p* = 0.3144).

**Figure 4.**
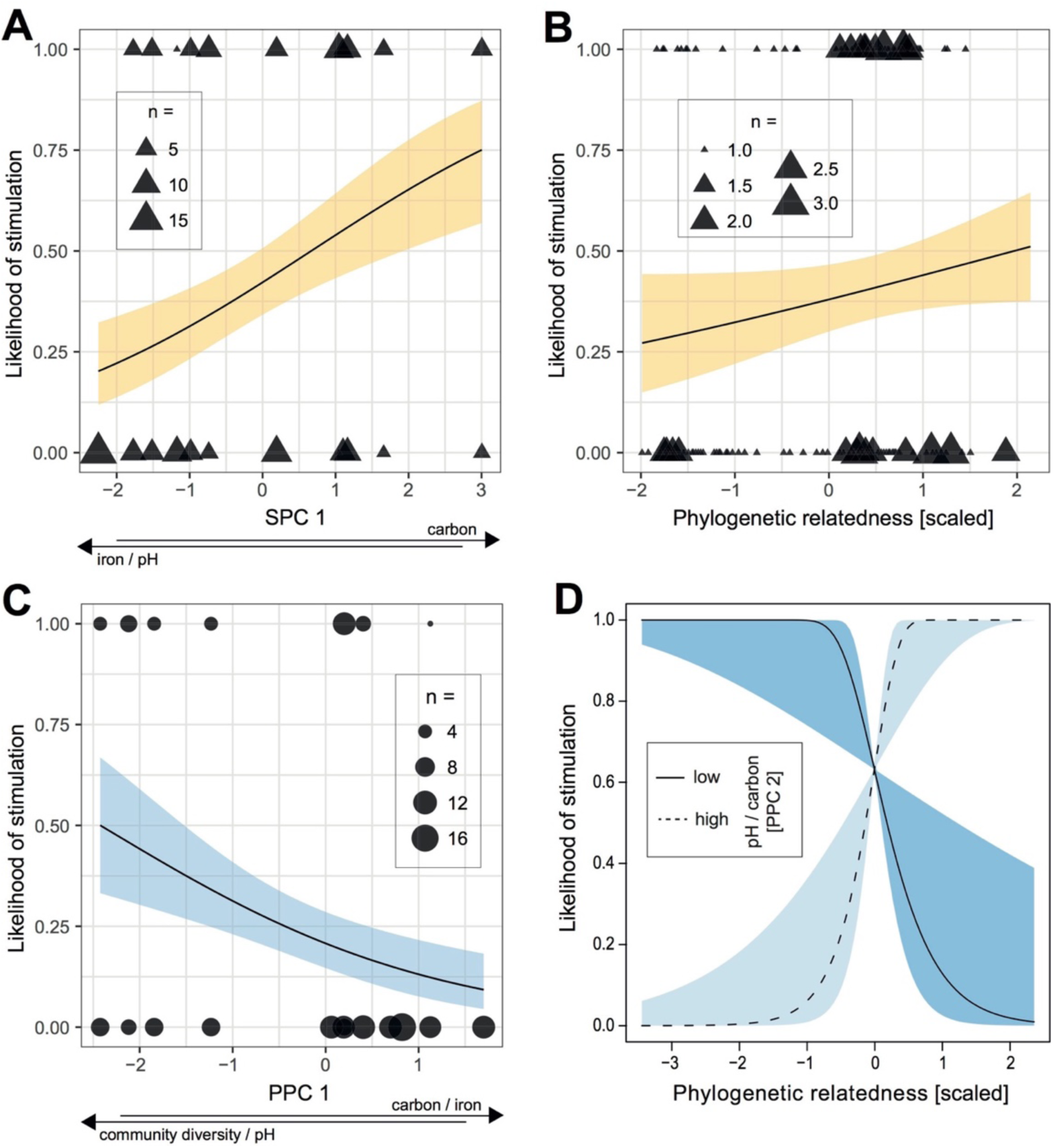
Correlations between parameters characterizing pyoverdine-mediated social interactions, environmental variables, and phylogenetic relatedness between strains. For soil non-producers, the likelihood to be stimulated by supernatants containing pyoverdine from a producer correlated positively with SPC1, i.e. it increased with increased carbon concentration, but reduced levels of pH and iron **(A)**; and phylogenetic relatedness between non-producer and producer based on *rpoD* identity **(B)**. For pond non-producers, the likelihood to be stimulated by producer supernatants correlated negatively with PPC1, i.e. it decreased with decreased pH and community diversity combined with increased carbon and iron concentrations **(C)**. In addition, there was a significant interaction between the *rpoD*-based phylogenetic relatedness between isolates and PPC2, whereby the likelihood of stimulation significantly increased with phylogenetic relatedness for higher values of pH and carbon (high PPC2), but decreased with phylogenetic relatedness for lower values of pH and carbon (low PPC2) **(D)**. Solid lines with shaded areas show significant correlations together with the 95% confidence band for soil (yellow) and pond (blue) communities. Data shown is based on 152 non-producer-supernatant combinations for soil, and 151 non-producer-supernatant combinations for pond.

For pond communities, we observed that the likelihood of stimulation negatively correlated with PPC1, suggesting that non-producers were most likely to be stimulated by heterologous pyoverdine(s) produced by other community members when originating from environments featuring low levels of carbon and iron together with high pH and community diversity (*z* = −2.13, *p* = 0.0333; Fig. 4C). Moreover, the likelihood of stimulation depended on an interaction between PPC2 and the phylogenetic relatedness between the supernatant donor and the recipient (interaction: *z* = 2.72, *p* = 0.0066; main effects: *rpoD* identity; *z* = 1.31, *p* = 0.1900; and PPC2; *z* = 0.65, *p* = 0.5132; Fig. 4D). In particular, a relatively high phylogenetic relatedness increased the likelihood of stimulation in communities featuring high levels of PPC2 (i.e. simultaneously high levels of pH and carbon), whereas this relationship was reversed in communities featuring low levels of PPC2 (Fig. 4D). By contrast, neither the PPCs nor *rpoD* identity correlated with the likelihood of inhibition (PPC1: *z* = 1.82, *p* = 0.0688; PPC2: *z* = −1.30, *p* = 0.1943; *rpoD* identity: *z* = −1.37, *p* = 0.1701).

## Discussion

The beauty of laboratory experiments in microbiology is that factors influencing bacterial physiology, behavior, and fitness can be investigated one at the time under controlled and replicable conditions. This approach contrasts with the situation bacteria typically face in nature, where environmental conditions often fluctuate rapidly in unpredictable manners, with multiple variables simultaneously influencing bacterial behavior and fitness. This raises the question whether the factors affecting bacterial behavior in vitro also play a role under natural conditions. Our study tackled this question by examining whether factors shown to influence an important bacterial trait in the laboratory, the production of siderophores used for iron-scavenging, also affect this behavior in natural communities. As a model system, we focused on the siderophore pyoverdine produced by *Pseudomonas* bacteria. Our investigations involving 930 *Pseudomonas* isolates, originating from 24 soil and 24 pond communities, yielded several novel insights (see Fig. 5 for a summary). First, we found that pH, concentrations of total iron and carbon, and community diversity, all shown to be important determinants of pyoverdine production in the laboratory, are indeed correlated with the level of pyoverdine produced by natural *Pseudomonas* isolates. Second, we observed that the same environmental variables also correlated with pyoverdine-mediated social interactions, measured by the extent to which secreted pyoverdine could promote or inhibit the growth of other members of the community. Third, we showed that the way these environmental variables correlated with pyoverdine production and social interactions differed fundamentally between soil and pond communities. Finally, our data suggest that trade-offs and interactions between environmental factors, which are typically ruled out in the laboratory, could be more predictive of bacterial behavior in nature than the main effect of a single factor.

**Figure 5.**
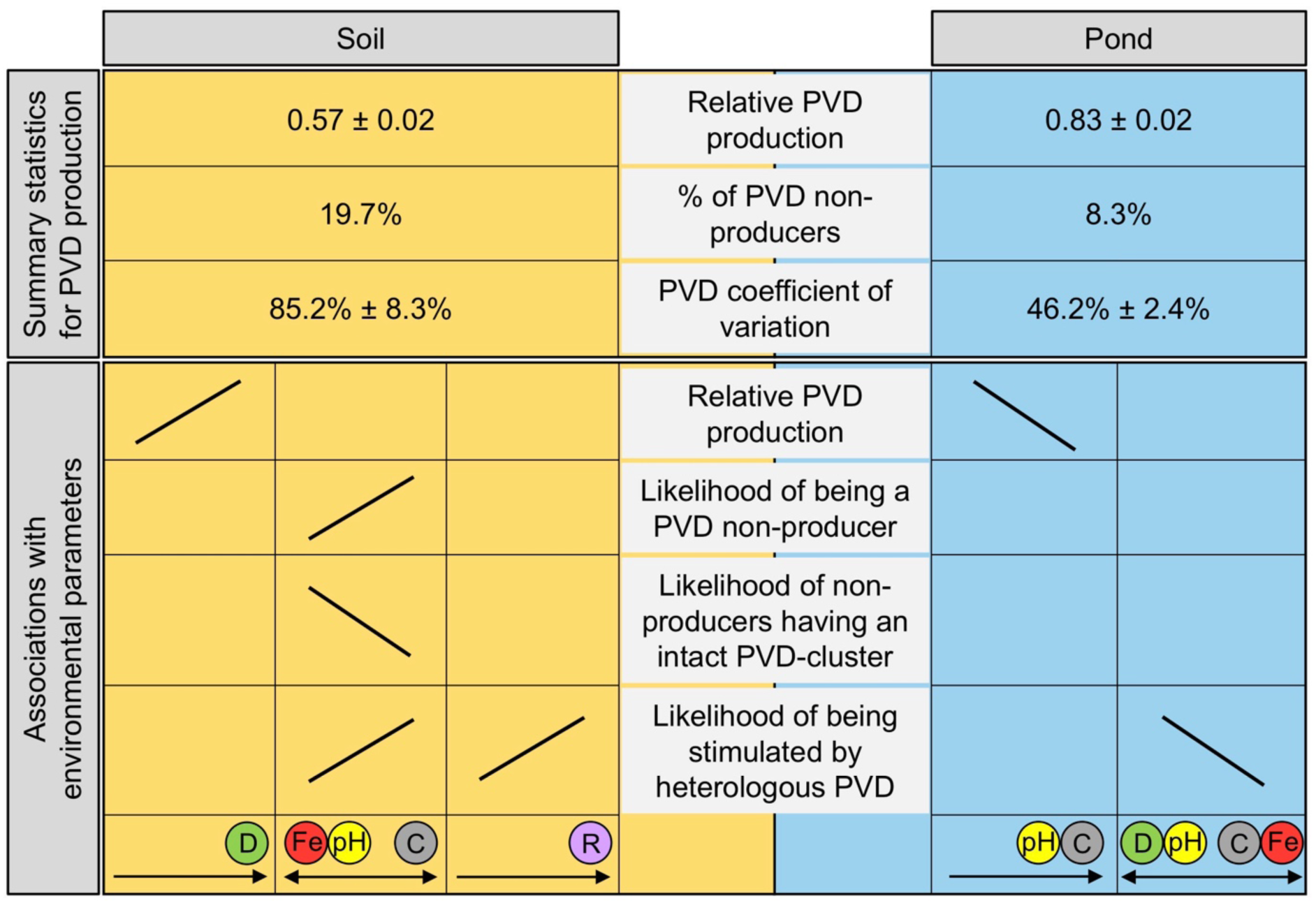
Overview of the main results on the determinants of pyoverdine (PVD) production in *Pseudomonas* soil and pond communities. The upper part of the scheme shows that: soil isolates produce on average less pyoverdine than pond isolates, there are more pyoverdine non-producers in soils than in ponds, and the coefficient of variation for pyoverdine production is higher in soil compared to pond communities. The lower part of the scheme indicates that there were more associations between environmental parameters and variables of pyoverdine production in soil than in pond communities. For isolates from soil communities, relative pyoverdine production was positively associated with community diversity (green circle), whereas variables of pyoverdine non-production all correlated with the environmental trade-off between iron concentration (red circle) and pH (yellow circle) versus carbon concentration (grey circle). Furthermore, the likelihood of non-producers being stimulated by heterologous pyoverdines increased with the relatedness between donor and recipient strain (purple circle). For isolates from pond communities, relative pyoverdine production was negatively associated with pH and carbon concentration. The likelihood of non-producers being stimulated by heterologous pyoverdines correlated with a complex trade-off between community diversity and pH versus carbon and iron concentration. For all isolates, pyoverdine production was scaled relative to laboratory reference strains (see Table S1). Variation in the environmental variables across soil and pond communities is shown in Fig. S3.

We observed that isolates from soil and pond communities differ substantially in their pyoverdine production profiles and in the way their production levels seem to be influenced by environmental factors. In particular, we found that: (a) soil isolates produced on average significantly less pyoverdine than pond isolates (Fig. 1A); (b) there were significantly more pyoverdine non-producers in soil than in pond (Fig. 1A; Table S2); and (c) the four environmental variables examined (community diversity, pH, total iron and carbon concentrations) were all associated with at least one aspect of pyoverdine production for soil isolates, whereas the pyoverdine production of pond isolates seemed to be much less affected by these variables (Fig. 2+5). Observation (a) can relatively easily be explained by the fact that diffusivity is higher and iron concentration orders of magnitude lower in ponds compared to soils (Fig. S3, Table S2). Although we need to be careful because not all the iron measured in our samples may actually be bioavailable, it seems that this element is more limited in ponds, such that higher siderophore investments are required for its scavenging. Observation (b) was more surprising, as we initially expected the opposite pattern, if non-producers evolve as cheaters, exploiting pyoverdines produced by others for iron scavenging. This is because the more diffusive nature of ponds combined with low iron availability should provide ample opportunities for pyoverdine exploitation. The opposite pattern observed here could suggest that there are other forces than cheating that select for non-production (see detailed discussion in next paragraph). Observation (c), meanwhile, could be the result of ponds being more open systems, where strain mixing is conceivably high due to the more diffusive and less structured environment, and where new strains can easily be introduced from the surrounding (terrestrial) habitats. Contrary to soils, where local adaptation has been observed (Bruce, West, *et al*., 2017), increased strain mixing and the potential source-sink dynamics between soil and pond habitats might even out local variation in community composition in ponds.

We propose three different mutually non-exclusive reasons for the evolution of pyoverdine non-producers in natural habitats. First, non-producers could evolve as cheaters exploiting the pyoverdine produced by others. In pond communities, this is a reasonable scenario because the prevailing conditions of high strain mixing, severe iron limitation and low carbon availability have been shown to promote cheating in laboratory experiments (Brockhurst *et al*., 2008; Kümmerli *et al*., 2009; Sexton and Schuster, 2017). Furthermore, aquatic bacteria often assemble on particles (Pernthaler, 2017), enabling non-producers to be close to producers, which is important for cheating (Cordero *et al*., 2012; Weigert and Kümmerli, 2017). In soil communities, on the other hand, cheaters could be favoured because this habitat sustains high cell densities, a factor previously shown to promote cheating (Ross-Gillespie *et al*., 2009; Scholz and Greenberg, 2015). While the complexity of natural habitats makes it difficult to assess the relative importance of these selection pressures, both habitats clearly have features that could select for cheating. Second, non-producers could evolve under conditions where pyoverdine is not stringently needed for iron acquisition. The idea that disuse drives pyoverdine loss is supported by our observation that the likelihood of being a non-producer was highest under conditions with highest predicted iron bioavailability (i.e. in soils with high carbon concentration in combination with low pH and iron concentration; Fig. 2B). Iron bioavailability is predicted to be highest under these conditions because iron solubility increases at low pH in soil, with the total iron content playing a minor role (Petruzzelli, 1989; Rieuwerts *et al*., 1998). Moreover, high carbon concentrations decrease the cost of pyoverdine production (Brockhurst *et al*., 2008; Sexton and Schuster, 2017), and can also increase the bioavailability of iron, via the metal-complexing properties of organic compounds, such as humic and fulvic acids (Lindsay, 1991; Hassler *et al*., 2009; Krachler *et al*., 2015). The hypothesis that pyoverdine is lost due to disuse is further supported by our observation that the frequency of non-producers, which have permanently lost the pyoverdine locus (i.e. *pvdL*-negative isolates), peaked exactly under the conditions of predicted highest iron bioavailability (Fig. 2D). Finally, non-producers could also evolve in the context of heavy metal detoxification. For instance, low pH in soils is known to increase the solubility of toxic heavy metals (Petruzzelli, 1989). Pyoverdine can bind many of those metals (usually with lower affinity than iron), and thereby contribute to the detoxification of the environment (Braud *et al*., 2009; Schalk *et al*., 2011). Since detoxification also constitutes a public good, it is possible that non-producers evolved as cheaters in a context of heavy metal detoxification (O’Brien *et al*., 2014; Hesse *et al*., 2018).

We now turn to our findings from the supernatant assay, where we fed pyoverdine-containing supernatants from producers to non-producers from the same community. For soil communities, we found that the likelihood of non-producers to be stimulated by supernatants increased under conditions of higher predicted iron availability (Fig. 4A) and when relatedness among strains was high (Fig. 4B). This could indicate that in relatively iron-replete soils and among closely related strains, the selection pressure for producers to evolve cheating-resistance mechanisms is reduced. Conversely, under stringent iron limitation (i.e. high pH combined with low carbon concentration) the likelihood of pyoverdine-mediated stimulation was lower, and inhibition increased. Under these conditions, selection thus seem to favour exclusive non-exploitable pyoverdines, for which non-producers lack uptake receptors (Smith *et al*., 2005; Lee *et al*., 2012). This is conceivable because cheating is thought to be especially harmful for producers when iron is scarce and carbon as a building block for pyoverdine is limited (Brockhurst *et al*., 2008; Kümmerli *et al*., 2009; Sexton and Schuster, 2017). Taken together, the patterns discovered here suggest that the role of pyoverdine changes with increasing iron availability: from a shareable, yet exploitable public good, to an inhibitory substance, locking iron away from competitors lacking a matching uptake receptor (Buyer and Leong, 1986; Butaitė *et al*., 2017; Niehus *et al*., 2017; Schiessl *et al*., 2017; Sexton *et al*., 2017).

For pond communities, we initially proposed that the openness of aquatic systems, characterized by increased strain influx from surrounding terrestrial habitats and increased strain mixing, reduce the potential for local strain adaptation to abiotic factors. These environmental properties are likely to also affect pyoverdine-mediated social interaction patterns. For instance, high levels of mixing could promote random interactions between pyoverdine producers and non-producers. This in turn could reduce the likelihood of non-producers to possess a matching receptor for the uptake of a specific pyoverdine, and thus lead to the high levels of pyoverdine-mediated growth inhibition observed in our pond communities (Fig. 3B). Nonetheless, we also found evidence for non-random interaction patterns, as the likelihood of non-producers to be stimulated by pyoverdine-containing supernatant from producers was highest in ponds with low predicted iron availability (high pH combined with low carbon and iron concentrations, but also high community diversity; Fig. 4C). This pattern opposes the one observed for soil non-producers (Fig. 4A), suggesting that selection for cheating and cheating resistance differ between the two habitat types. This notion is also supported by the occurrence of complex interactions between inter-strain relatedness and the environmental variables in ponds (Fig. 4D), but not in soils.

In conclusion, our study reveals that connecting laboratory to field studies is a challenging task for microbiologists. While laboratory studies are typically carried out with isogenic strains and engineered single-gene knockout mutants grown under controlled conditions, field studies, like ours, face an enormous strain diversity, and a plethora of environmental factors that all simultaneously vary and influence bacterial behavior. Despite this complexity, we managed to obtain first insights into the importance of siderophore-based iron acquisition and competition in natural communities of pseudomonads. In particular, we found evidence that pyoverdine production levels of natural isolates and the ability to use heterologous pyoverdine are simultaneously influenced by multiple environmental variables, including pH, iron and carbon concentrations, with patterns clearly differing between soil and pond habitats. But our study also faces some limitations, as it focusses solely on pyoverdine and pseudomonads, thereby ignoring the fact that stealing of heterologous siderophores can occur across the genus boundary, and involve different siderophore systems (D’Onofrio *et al*., 2010; Galet *et al*., 2015). Moreover, while our standardized laboratory assay allowed us to assess the strains’ genetic potential to make pyoverdine, it does not inform us on how much pyoverdine these strains actually make in their native habitat, where strains might flexibly adjust their pyoverdine production to short-term variation in resource availability and/or the presence of competitors. These considerations show that there is still a long way until we will fully understand how complex microbial communities operate. For all these reasons, we advocate the need for further studies that embrace the complexity of natural systems, in order to unravel how microbial behavior and strain interactions shape community composition and functioning.

## Experimental Procedures

### Sampling and isolation of pseudomonads

We sampled soil cores and water samples at three different sites in late May 2013 (water samples) and early July 2013 (soil samples). While one site was situated on the Irchel campus of the University of Zurich (47.40° N, 8.54° E), the other two sites were located at Seleger Moor Park (47.25° N, 8.51° E), Switzerland. At each site, we collected eight soil cores and eight water samples for each habitat type (soil or pond), resulting in a total of 48 samples. We followed a rectangular collection scheme, whereby each soil core/water sample had one close (50 cm), two intermediate (5 m) and four distant (50 m) neighboring samples. This was done to examine the effect of geographical distance on community structure, which is, however, not the focus of this study. The a priori reasoning for sampling at these different sites was that we expected ecological parameters to differ quite substantially between the habitats in a city park and more natural environments. From the 24 soil cores and 24 pond water samples, we isolated 952 strains (target: 20 per sample) on a medium selective for fluorescent pseudomonads, and preserved the isolates as freezer stocks as described in detail elsewhere (Butaitė *et al*., 2017). We provided each isolate with a unique identification code, consisting of a site ID (‘s1 = Seleger Moor soil#1’, ‘s2 = Seleger Moor soil#2’, ‘s3 = Irchel soil’, ‘1 = Seleger Moor pond#1’, ‘2 = Seleger Moor pond#2’, ‘3 = Irchel pond’), a community ID (small letters ‘a’ to ‘h for soil communities; capital letters ‘A’ to ‘H’ for pond communities) and an isolate number (1 to 20).

### Measuring environmental parameters

We measured the pH of soil and pond samples using the Atago portable pH meter DPH-2 (for field measurements) and the Metrohm 744 pH meter (for laboratory measurements). The pH of pond samples was first measured in the laboratory and then later confirmed by measurements directly in the field. Prior to soil pH measurements, the soil samples were suspended 1:5 (w/v) in 0.01 M CaCl_2_ solution, shaken for 1 h, and left to sediment.

We measured the percentage of total carbon in soil and pond samples with a CHN (Leco Truspec micro; Leco Instruments, USA) and a TOC analyser (Dimatoc 2000; Dimatec, Germany), respectively. Measurements were carried out by staff operating the specific in-house service at the University of Zurich. Prior to the measurements, we first dried soil samples for two days at 40 °C, then grounded them to a homogenous powder and dried them again at 40 °C overnight.

To estimate the iron concentration of the sampled soils and ponds, we used the ICP-OES methodology (Vista-MPX, Varian, USA). For soils, we first dried and homogenized the plant-free samples (plant material was removed using a 0.6 mm stainless steel sieve). Soil samples were then digested with HCl and HNO_3_ for 90 min at 120 °C, diluted in Nanopure water and filtered. The pond samples were directly filtered using a 0.2 µm filter and then acidified with 65% HNO_3_ to pH < 3. To make iron concentrations comparable between soil (µg/g) and pond (µg/L) samples, we converted pond concentrations to µg/g by assuming that 1 ml = 1 g.

### *rpoD* amplification and sequencing

To verify that the 952 isolates are indeed *Pseudomonas*, we PCR amplified and sequenced a part of the *rpoD* gene for 930 isolates (PCR or sequencing failed for 22 isolates, which were excluded from further experiments). This housekeeping gene is commonly used for phylogenetic affiliation of pseudomonads (Mulet *et al*., 2009; Ghyselinck *et al*., 2013). PCR mixtures were prepared, PCR reactions were carried out and the products were sequenced using PsEG30F and PsEG790R primers (Mulet *et al*., 2009) as described elsewhere (Butaitė *et al*., 2017).

### Community diversity and phylogenetic relatedness

A codon-aware nucleotide alignment of *rpoD* was generated using local TranslatorX v1.1 (Abascal *et al*., 2010) with the MAFFT v7.271 (Katoh and Standley, 2013) aligner. We manually curated and trimmed the alignment at both ends resulting in a high-quality alignment of 908 sequences (including 21 reference strains; as described elsewhere (Butaitė *et al*., 2017) over 609 nucleotides. The phylogenetic tree was inferred by RAxML v7.0.4 (Stamatakis, 2006) using the General Time Reversible (GTR) + G model with 100 bootstraps. We calculated phylogenetic diversity within communities based on the cophenetic function from the ape package in R software and normalized values by the number of tips. For some analyses, we calculated the relatedness between strains by carrying out pairwise alignments of *rpoD* sequences using the ‘water’ application from EMBOSS (Rice *et al*., 2000).

### *pvdL* amplification

We used the presence of the *pvdL* gene as a proxy for the presence of the pyoverdine biosynthesis locus in all 130 pyoverdine non-producers. *pvdL* is involved in the biosynthesis of the pyoverdine chromophore, and is the most conserved part of the locus across different pseudomonads (Ravel and Cornelis, 2003). To identify aconserved region of this gene, we aligned the *pvdL* sequences (retrieved from GenBank) of seven pseudomonads: *P. aeruginosa* PAO1, *P. chlororaphis* O6, *P. chlororaphis subsp.aureofaciens* 30–84, *P. fluorescens* A506, *P. fluorescens* SS101, *P. protegens* Pf-5, *P. syringae pv.syringae* B64. We then designed the following primers to amplify a conserved region in the isolated non-producers (pvdL_fw: CATGATGAGCAACCACCACATC, pvdL_rv: CGCTGGTCGTAGGACAGGTG; product size: 827 bp). We prepared PCR mixtures as for the *rpoD* gene amplification (Butaitė *et al*., 2017), just this time we used a ‘Hot Start’ version of the Master Mix. Bacterial biomass was taken from fresh cultures grown in liquid LB medium (1 µl). We used the following PCR conditions: denaturation at 94.5 °C for 5 min; 30 cycles of amplification (1 min denaturation at 94 °C, 1 min primer annealing at 57 °C, and 1 min primer extension at 68 °C); final elongation at 68 °C for 10 min. As a positive control, we used 72 pyoverdine producers (pond and soil, 36 each). We considered an isolate as *pvdL*-positive when the PCR yielded a product of the expected size. Among the 72 pyoverdine producers (which should all have *pvdL*), we found seven to be negative for *pvdL*. We thus estimate the rate of false-negatives to be 9.7% (i.e. the risk to falsely identify isolates as *pvdL*-negatives).

### Measurement of growth and pyoverdine production levels

To evaluate whether and to what extent the isolates can produce pyoverdine, we grew them under iron-limited conditions and measured their pyoverdine production levels. We first grew the isolates in 150 µl LB in 96-well plates overnight (16 - 18 h) static at room temperature. We then transferred 2 µl of overnight cultures to 200 µl iron-limited casamino acids medium (CAA containing 5 g casamino acids, 1.18 g K_2_HPO_4_·3H_2_O, 0.25 g MgSO_4_·7H_2_O per liter) supplemented with 25 mM HEPES buffer, 20 mM NaHCO_3_ and 100 µg/ml of the strong iron chelator human apo-transferrin in a 96-well plate. All chemicals were purchased from Sigma-Aldrich, Switzerland.

Each plate had isolates from one community in triplicates and eight reference strains known to produce pyoverdine (see Table S1 for a description of these strains). After 18 h of incubation at room temperature, we measured growth (optical density OD at 600 nm) and pyoverdine production levels (relative fluorescence units, RFU, with excitation: 400 nm and emission: 460 nm) with an Infinite M200 Pro microplate reader (Tecan Group Ltd., Switzerland). We then scaled the growth and relative pyoverdine production values for each isolate by dividing their OD_600_ and RFU by the average respective OD_600_ and RFU of the reference strains.

We also carried out a control experiment to verify that it is indeed iron limitation that induces the observed growth and pyoverdine production patterns, and not a specific effect of transferrin as an iron chelator. For this, we repeated the growth assay with the synthetic iron chelator 2,2’-dipyridyl (200 µM). The growth assay was carried out for a subset of soil (n = 100: 28 pyoverdine non-producers and 72 producers) and pond isolates (n = 99: 27 non-producers and 72 producers), from 12 soil and 12 pond communities, in three replicates. The assay yielded qualitative very similar results for both iron chelators (Fig. S5). Moreover, in an earlier study we could demonstrate that the environmental isolates (*n* = 315) could use CAA as a carbon source, demonstrating that the poor growth of non-producers is due to their inability to produce pyoverdine, and not because they are unable to grow in CAA (Butaitė *et al*., 2017).

### Supernatant assay

To quantify the effect that pyoverdine-containing supernatants from producers has on non-producers, we set up a supernatant growth assay using a subset of isolates and communities (24 communities in total: four communities per site and habitat type). For each community, we initially aimed at selecting six pyoverdine producers that (a) grew better than the corresponding non-producers under iron-limited condition; and (b) differed in their *rpoD* sequence, and thus represented phylogenetically different strains. These criteria were met for 22 communities. For one soil community (s2e), we were limited to four producers satisfying (a), and for two communities (3E and s2e), we had to include two pairs of producers with identical *rpoD* sequences. Overall, we used 72 pond and 70 soil producers. For each community, we further randomly selected up to three non-producers (upon availability). In total, we had 26 and 27 non-producers (relative pyoverdine production < 5% relative to the eight reference pyoverdine producers listed in Table S1) from soil and pond communities, respectively. This resulted in 152 soil non-producer-supernatant and 151 pond non-producer-supernatant combinations.

To generate pyoverdine-containing supernatants, we first grew all producers in 200 µl LB medium in 96-well plates overnight at 25 °C. Then, 20 µl of overnight cultures were added to 2 ml of CAA with 200 µM 2,2’-dipyridyl in 24-well plates and incubated static at 25 °C for about 22 h. We centrifuged cultures for 10 min at 3,500 rpm (Eppendorf Centrifuge 5804R) and then transferred 900 µl of supernatants to PALL AcroPrep Advance 96-well 1 ml filterplates (with 0.2 µm supor membrane), attached to an autoclaved 1.2 ml 96-well PCR plate (VWR). We centrifuged the assemblies of filter and collection plates at 2,500 rpm for 15 min. The collection plates with sterile supernatants and blank medium were sealed with Greiner SILVERseal and kept at −20 °C.

Afterwards we grew the non-producers in 200 µl of LB in 96-well plates overnight static at 25 °C. We then added 2 µl of non-producer cultures to CAA with 200 µM 2,2’-dipyridyl supplemented with 20 µl of (a) producer supernatant or (b) CAA with 200 µM 2,2’-dipyridyl that went through the same treatment as supernatants, i.e. filtering and freezing. Total culturing volume was 200 µl. Each treatment was set up in four replicates. Plates were incubated static for 17 h at 25 °C. The final OD_600_ of the cultures was measured using the Tecan microplate reader. We considered a supernatant effect as ‘stimulation’ or ‘inhibition’ when non-producers grew significantly better or worse, respectively, with a supernatant than without, based on a Wilcoxon test.

### Statistical analysis

We used linear mixed-effects (LMM) and linear generalized mixed-effects (GLMM) models for statistical data analysis. Since neighboring communities (50 cm apart) might be more similar to each other than to the more distant communities from the same site, we included both ‘close neighbors’ and ‘community’ (nested within ‘close neighbors’) as random effects into our models. Whenever appropriate, we used natural log-transformations to meet the assumption of normally distributed residuals. For the supernatant assays, we used binomial models to calculate the likelihood of supernatant receivers in a given community to be stimulated or inhibited. Note that the likelihood of stimulation/inhibition can also be interpreted as the proportion of ‘stimulatory’ or ‘inhibitory’ interactions observed within a community. All statistical analyses were carried out using the R 3.1.2 statistics software (www.r-project.org).

The environmental variables were often highly correlated (Table S3), which led to high collinearities in statistical models (variance inflation factor (vif) >> 10). We thus conducted principal component analyses (PCAs) to obtain non-correlated principal components (PCs) reflecting single or combinations of different environmental variables (entered after centering and scaling to unit variance, respectively). We performed two separate PCAs for soil and pond communities, because the overall differences in pH and the total concentrations of iron and carbon (Table S2), together with differences in the signs of some of the correlations among environmental variables in the soil and pond (Table S3), also lead to high collinearity (variance inflation factor >> 10) in models deploying PCs based on the whole dataset. We thus used the PCs resulting from separate PCAs for soil and pond (Table 1) to analyse their effects on pyoverdine production and social interactions.

## Acknowledgments

We dedicate this article to Prof. Helmut Brandl, who inspired and advised us on this project, but sadly passed away in 2017. We thank Roland Dünner for permission to sample at Seleger Moor park, René Husi for carbon concentration measurements, and Prof. Rainer Schulin and Björn Studer from ETH Zurich for iron concentration measurements at their facility. This work was funded by the Swiss National Science Foundation (grants no. 139164 and 165835 to RK) and the Forschungskredit of the University of Zurich (FK-15–082 to EB and FK-17–111 to JK). The authors declare no conflict of interest.

## Author contribution

EB and RK designed the study, EB isolated the strains and performed the experiments, EB, JK, SW and RK analyzed the data and wrote the paper.

## References

Abascal, F., Zardoya, R., and Telford, M.J. (2010) TranslatorX: multiple alignment of nucleotide sequences guided by amino acid translations. Nucleic Acids Res. 38: W7–W13.

Andrews, S.C., Robinson, A.K., and Rodríguez-Quiñones, F. (2003) Bacterial iron homeostasis. FEMS Microbiol. Rev. 27: 215–237.

Boyd, P.W. and Ellwood, M.J. (2010) The biogeochemical cycle of iron in the ocean. Nat. Geosci. 3: 675–682.

Braud, A., Hoegy, F., Jezequel, K., Lebeau, T., and Schalk, I.J. (2009) New insights into the metal specificity of the *Pseudomonas aeruginosa* pyoverdine-iron uptake pathway. Environ. Microbiol. 11: 1079–1091.

Brockhurst, M.A., Buckling, A., Racey, D., and Gardner, A. (2008) Resource supply and the evolution of public-goods cooperation in bacteria. BMC Biol. 6: 20.

Bruce, J.B., Cooper, G.A., Chabas, H., West, S.A., and Griffin, A.S. (2017) Cheating and resistance to cheating in natural populations of the bacterium *Pseudomonas fluorescens*. Evolution 71: 2484–2495.

Bruce, J.B., West, S.A., and Griffin, A.S. (2017) Bacteriocins and the assembly of natural *Pseudomonas fluorescens* populations. J. Evol. Biol. 30: 352–360.

Butaitė, E., Baumgartner, M., Wyder, S., and Kümmerli, R. (2017) Siderophore cheating and cheating resistance shape competition for iron in soil and freshwater *Pseudomonas* communities. Nat. Commun. 8: 414.

Buyer, J. and Leong, J. (1986) Iron transport-mediated antagonism between plant growth-promoting and plant-deleterious *Pseudomonas* strains. J. Biol. Chem. 261: 791–794.

Cézard, C., Farvacques, N., and Sonnet, P. (2015) Chemistry and biology of pyoverdines, *Pseudomonas* primary siderophores. Curr. Med. Chem. 22: 165–186.

Cordero, O.X., Ventouras, L.-A., DeLong, E.F., and Polz, M.F. (2012) Public good dynamics drive evolution of iron acquisition strategies in natural bacterioplankton populations. Proc. Natl. Acad. Sci. U. S. A. 109: 20059–64.

Cornelis, P. (2010) Iron uptake and metabolism in pseudomonads. Appl. Microbiol. Biotechnol. 86: 1637–1645.

D’Onofrio, A., Crawford, J.M., Stewart, E.J., Witt, K., Gavrish, E., Epstein, S., et al. (2010) Siderophores from neighboring organisms promote the growth of uncultured bacteria. Chem. Biol. 17: 254–264.

Dumas, Z., Ross-Gillespie, A., and Kümmerli, R. (2013) Switching between apparently redundant iron-uptake mechanisms benefits bacteria in changeable environments. Proc. Biol. Sci. 280: 20131055.

Frawley, E.R. and Fang, F.C. (2014) The ins and outs of bacterial iron metabolism. Mol. Microbiol. 93: 609–616.

Galet, J., Deveau, A., Hôtel, L., Frey-Klett, P., Leblond, P., and Aigle, B. (2015) *Pseudomonas fluorescens* pirates both ferrioxamine and ferricoelichelin siderophores from *Streptomyces ambofaciens*. Appl. Environ. Microbiol. 81: 3132–3141.

Ghyselinck, J., Coorevits, A., Van Landschoot, A., Samyn, E., Heylen, K., and De Vos, P. (2013) An *rpoD* gene sequence based evaluation of cultured *Pseudomonas* diversity on different growth media. Microbiology 159: 2097–108.

Griffin, A.S., West, S.A., and Buckling, A. (2004) Cooperation and competition in pathogenic bacteria. Nature 430: 1024–1027.

Harrison, F., Paul, J., Massey, R.C., and Buckling, A. (2008) Interspecific competition and siderophore-mediated cooperation in *Pseudomonas aeruginosa*. ISME J. 2: 49–55.

Hassler, C.S., Havens, S.M., Bullerjahn, G.S., McKay, R.M.L., and Twissa, M.R. (2009) An evaluation of iron bioavailability and speciation in western Lake Superior with the use of combined physical, chemical, and biological assessment. Limnol. Oceanogr. 54: 987–1001.

Hesse, E., O’Brien, S., Tromas, N., Bayer, F., Luján, A.M., van Veen, E.M., et al. (2018) Ecological selection of siderophore-producing microbial taxa in response to heavy metal contamination. Ecol. Lett. 21: 117–127.

Hider, R.C. and Kong, X. (2010) Chemistry and biology of siderophores. Nat. Prod. Rep. 27: 637.

Inglis, R.F., Biernaskie, J.M., Gardner, A., and Kümmerli, R. (2016) Presence of a loner strain maintains cooperation and diversity in well-mixed bacterial communities. Proc. Biol. Sci. 283: 20152682.

Jiricny, N., Diggle, S.P., West, S.A., Evans, B.A., Ballantyne, G., Ross-Gillespie, A., and Griffin, A.S. (2010) Fitness correlates with the extent of cheating in a bacterium. J. Evol. Biol. 23: 738–747.

Jolliffe, I. T. (1972) Discarding variables in a principal component analysis. I: artificial data. J. R. Stat. Soc. Series A. 21: 160–173.

Joshi, F., Archana, G., and Desai, A. (2006) Siderophore cross-utilization amongst rhizospheric bacteria and the role of their differential affinities for Fe3+ on growth stimulation under iron-limited conditions. Curr. Microbiol. 53: 141–147.

Katoh, K. and Standley, D.M. (2013) MAFFT multiple sequence alignment software version 7: improvements in performance and usability. Mol. Biol. Evol. 30: 772–80.

Krachler, R., Krachler, R.F., Wallner, G., Hann, S., Laux, M., Cervantes Recalde, M.F., et al. (2015) River-derived humic substances as iron chelators in seawater. Mar. Chem. 174: 85–93.

Kümmerli, R., Jiricny, N., Clarke, L.S., West, S.A., and Griffin, A.S. (2009) Phenotypic plasticity of a cooperative behaviour in bacteria. J. Evol. Biol. 22: 589–598.

Lee, W., van Baalen, M., and Jansen, V.A.A. (2012) An evolutionary mechanism for diversity in siderophore-producing bacteria. Ecol. Lett. 15: 119–125.

Leinweber, A., Inglis, R.F., and Kümmerli, R. (2017) Cheating fosters species co-existence in well-mixed bacterial communities. ISME J. 11: 1179–1188.

Leinweber, A., Weigert, M., and Kümmerli, R. (2018) The bacterium *Pseudomonas aeruginosa* senses and gradually responds to inter-specific competition for iron. Evolution (in press).

Lindsay, W.L. (1991) Iron oxide solubilization by organic matter and its effect on iron availability. Plant Soil 130: 27–34.

Loper, J.E. and Henkels, M.D. (1997) Availability of iron to *Pseudomonas fluorescens* in rhizosphere and bulk soil evaluated with an ice nucleation reporter gene. Appl. Environ. Microbiol. 63: 99–105.

Loper, J.E. and Henkels, M.D. (1999) Utilization of heterologous siderophores enhances levels of iron available to *Pseudomonas putida* in the rhizosphere. Appl. Environ. Microbiol. 65: 5357–63.

Meyer, J.-M., Gruffaz, C., Raharinosy, V., Bezverbnaya, I., Schäfer, M., and Budzikiewicz, H. (2008) Siderotyping of fluorescent *Pseudomonas*: molecular mass determination by mass spectrometry as a powerful pyoverdine siderotyping method. BioMetals 21: 259–271.

Miethke, M. and Marahiel, M. a (2007) Siderophore-based iron acquisition and pathogen control. Microbiol. Mol. Biol. Rev. 71: 413–451.

Mulet, M., Bennasar, A., Lalucat, J., and García-Valdés, E. (2009) An *rpoD*-based PCR procedure for the identification of *Pseudomonas* species and for their detection in environmental samples. Mol. Cell. Probes 23: 140–7.

Neilands, J.B. (1981) Iron absorption and transport in microorganisms. Annu. Rev. Nutr. 1: 27–46.

Niehus, R., Picot, A., Oliveira, N.M., Mitri, S., and Foster, K.R. (2017) The evolution of siderophore production as a competitive trait. Evolution 71: 1443–1455.

O’Brien, S., Hodgson, D.J., and Buckling, A. (2014) Social evolution of toxic metal bioremediation in *Pseudomonas aeruginosa*. Proc. Biol. Sci. 281: 20140858.

Parrow, N.L., Fleming, R.E., and Minnick, M.F. (2013) Sequestration and scavenging of iron in infection. Infect. Immun. 81: 3503–14.

Pernthaler, J. (2017) Competition and niche separation of pelagic bacteria in freshwater habitats. Environ. Microbiol. 19: 2133–2150.

Petruzzelli, G. (1989) Recycling wastes in agriculture: heavy metal bioavailability. Agric. Ecosyst. Environ. 27: 493–503.

Ravel, J. and Cornelis, P. (2003) Genomics of pyoverdine-mediated iron uptake in pseudomonads. Trends Microbiol. 11: 195–200.

Rice, P., Longden, I., and Bleasby, A. (2000) EMBOSS: the European Molecular Biology Open Software Suite. Trends Genet. 16: 276–7.

Rieuwerts, J.S., Thornton, I., Farago, M.E., and Ashmore, M.R. (1998) Factors influencing metal bioavailability in soils: preliminary investigations for the development of a critical loads approach for metals. Chem. Speciat. Bioavailab. 10: 61–75.

Ross-Gillespie, A., Gardner, A., Buckling, A., West, S.A., and Griffin, A.S. (2009) Density dependence and cooperation: Theory and a test with bacteria. Evolution 63: 2315–2325.

Schalk, I.J., Hannauer, M., and Braud, A. (2011) New roles for bacterial siderophores in metal transport and tolerance. Environ. Microbiol. 13: 2844–2854.

Schiessl, K.T., Janssen, E.M.-L., Kraemer, S.M., McNeill, K., and Ackermann, M. (2017) Magnitude and mechanism of siderophore-mediated competition at low iron solubility in the *Pseudomonas aeruginosa* pyochelin system. Front. Microbiol. 8: 1964.

Scholz, R.L. and Greenberg, E.P. (2015) Sociality in *Escherichia coli*: enterochelin is a private good at low cell density and can be shared at high cell density. J. Bacteriol. 197: 2122–8.

Sexton, D.J., Glover, R.C., Loper, J.E., and Schuster, M. (2017) *Pseudomonas protegens* Pf-5 favors self-produced siderophore over free-loading in interspecies competition for iron. Environ. Microbiol. 19: 3514–3525.

Sexton, D.J. and Schuster, M. (2017) Nutrient limitation determines the fitness of cheaters in bacterial siderophore cooperation. Nat. Commun. 8: 230.

Smith, E.E., Sims, E.H., Spencer, D.H., Kaul, R., and Olson, M. V. (2005) Evidence for diversifying selection at the pyoverdine locus of *Pseudomonas aeruginosa*. J. Bacteriol. 187: 2138–2147.

Stamatakis, A. (2006) RAxML-VI-HPC: maximum likelihood-based phylogenetic analyses with thousands of taxa and mixed models. Bioinformatics 22: 2688–2690.

Tiburzi, F., Imperi, F., and Visca, P. (2008) Intracellular levels and activity of PvdS, the major iron starvation sigma factor of *Pseudomonas aeruginosa*. Mol. Microbiol. 67: 213–227.

Traxler, M.F., Seyedsayamdost, M.R., Clardy, J., and Kolter, R. (2012) Interspecies modulation of bacterial development through iron competition and siderophore piracy. Mol. Microbiol. 86: 628–44.

Weigert, M. and Kümmerli, R. (2017) The physical boundaries of public goods cooperation between surface-attached bacterial cells. Proc. Biol. Sci. 284: 20170631.

West, S.A., Griffin, A.S., Gardner, A., and Diggle, S.P. (2006) Social evolution theory for microorganisms. Nat. Rev. Microbiol. 4: 597–607.

